# Fork in the road: how self-confidence about terrain influences gaze behaviour and path choice

**DOI:** 10.1101/2023.06.29.547105

**Authors:** Vinicius da Eira Silva, Daniel S. Marigold

## Abstract

Decisions about where to move occur throughout the day and are essential to life. Different movements may present different challenges and affect the likelihood of achieving a goal. Certain choices may have unintended consequences, some of which may cause harm and bias the decision. Movement decisions rely on a person gathering necessary visual information via shifts in gaze. Here we sought to understand what influences this information-seeking gaze behaviour. We had participants choose between walking across one of two paths that consisted of images of terrain commonly found in either hiking or urban environments. We manipulated the number and type of terrain of each path. We recorded gaze behaviour during the approach to the two paths and had participants rate their self-confidence about walking across each type of terrain as though it was real. Participants did not direct gaze to the path with greater visual information, regardless of how we quantified information. Rather, we show that a person’s self-confidence about their motor abilities predicts how they visually explore the environment with their eyes as well as their choice of action. The greater the self-confidence about walking across one path, the more they directed gaze to it, and the more likely they chose to walk across it. Overall, our results reveal a key role of a cognitive factor (self-confidence) in decision-making during a natural, movement-related behaviour.

## INTRODUCTION

Decisions require information about available choices. In complex environments, like a busy street, shopping mall, or hiking trail, there is a multitude of information that competes for attention and can collectively inform the eventual choice of action. Shifts in gaze allow a person to extract timely, high-resolution visual information necessary to make a goal-directed decision. But what drives the decision of where and for how long to direct gaze in these natural behaviours? And how does this information affect the decision about how to act?

In goal-directed, natural behaviours, gaze is directed to task-relevant features of the environment (Land et al. 1999; Marigold and Patla 2007; Rothkopf et al. 2007). More specifically, gaze is drawn to areas of uncertainty, thus allowing for increased information gain (Daddaoua et al. 2016; Domínguez-Zamora et al. 2018; Hayhoe 2017; Sprague and Ballard 2003; Sprague et al. 2007; Sullivan et al. 2012; Tong et al. 2017). However, people can exhibit different preferences for information gain, which predicts their subsequent choice of action (Domínguez-Zamora and Marigold 2021). When decisions are made during movement, such as walking or driving, there is limited time to gather information, and the information available may change during the decision process. Consequently, this requires a trade-off between exploring several areas with gaze and fixating on a restricted area of the environment. The decision to acquire further information may depend on a person’s confidence in the information they have currently collected. In support, previous work shows that confidence can influence further information-seeking behaviour, and that decisions are sensitive to the cost of obtaining the additional information (Boldt et al. 2019; Desender et al. 2018, 2019; Schulz et al. 2023).

Most decision-making studies define confidence in terms of the probability (or certainty) of being correct (e.g., Kiani & Shadlen 2009, Kiani et al 2014, Desender et al. 2018, Pouget et al. 2016). These studies typically use perceptual decision-making tasks to determine how being confident about a given choice relates to decision making. For example, Kiani et al (2014) demonstrated that confidence in making the correct choice inversely correlated with the decision’s reaction time, and positively correlated with the coherence of the presented sensory evidence in the task. However, not all decisions have a correct or incorrect outcome; in motor behaviour, each choice may achieve the same (or very similar) goal but in different ways. For example, when walking, cycling, or driving to get home, different routes allow you to reach your destination. Certain action choices may also have undesirable consequences. With walking, this may include a loss of balance and potential injury, getting lost, and/or expending greater energy due to the added distance to travel. The decision of which action to choose may depend, in part, on the perception of (or trust in) one’s abilities, or self-confidence. For instance, it is reasonable to expect that a person will favour an action they perceive poses less danger or is easier to handle. If gaze serves to gather relevant information about action alternatives, and a person can use this information to evaluate their confidence in performing each action, then might self-confidence affect gaze decisions in addition to action choice?

To address this question, we had participants decide between walking across one of two paths that we projected on the ground. Both paths had images of terrain commonly found in either hiking or urban settings. Because of the role of gaze in gaining information, we designed two protocols that varied in the information available in each walking path. In one, the two paths had a different number of terrain patches. In the other, both paths had an equal number of terrain patches but differed in terms of the information contained within the visual images. If information-seeking alone explains gaze behaviour, we would expect a greater number of fixations and/or longer gaze time on the path with greater information. However, we propose that self-confidence about walking across a path modifies information-seeking behaviour. Thus, we tested the hypothesis that self-confidence about walking across a path affects where and for how long gaze is directed to each path in addition to the choice of which path to take. We had participants rate their confidence in walking across each type of terrain as though they had to step on it in real life. We show that a simple information-seeking perspective is insufficient to explain gaze patterns when choosing between walking paths; participants did not direct gaze to the path with greater information, regardless of how we quantified information. Rather, we show that (1) less confidence on a uniform (single) terrain path associated with a greater number of fixations on a non-uniform (three) terrain path, and (2) with two non-uniform paths, greater self-confidence with the path eventually chosen, relative to the alternative, associated with a greater number of fixations and gaze time on it during the approach. These results support the notion that a person’s perception of their abilities impacts gaze decisions.

## RESULTS

Participants (n = 16) performed a visually guided walking paradigm that required them to choose and walk across one of two paths that we projected on the ground and had images of terrain found in either hiking or urban settings. We used images of terrain to provide greater control over the design of the paths and because they eliminate energetic and stability costs associated with walking across real terrain, which are known to affect gaze (Domínguez-Zamora and Marigold 2019, 2021; Matthis et al. 2018). Participants completed two different protocols (see Fig. 1), the order of which was counterbalanced. In one protocol, referred to as 1vs3, we used eight different environments (four hiking and four urban), where one path consisted of one type of terrain (uniform path) and one path consisted of three types of terrain (non-uniform path). In the other protocol, referred to as 3vs3, we used four different environments (two hiking and two urban), where each path consisted of three types of terrain. Participants started walking from approximately 1.5 m from the projected environment. We recorded gaze position using a mobile eye tracker and analyzed gaze data prior to and while walking until the participant reached the fork in the paths (i.e., approach phase). We had each participant rate their confidence level of each terrain based on how confident (or certain) they were about walking across it without losing balance, as though they had to step on it in real life outside.

**Figure 1:**
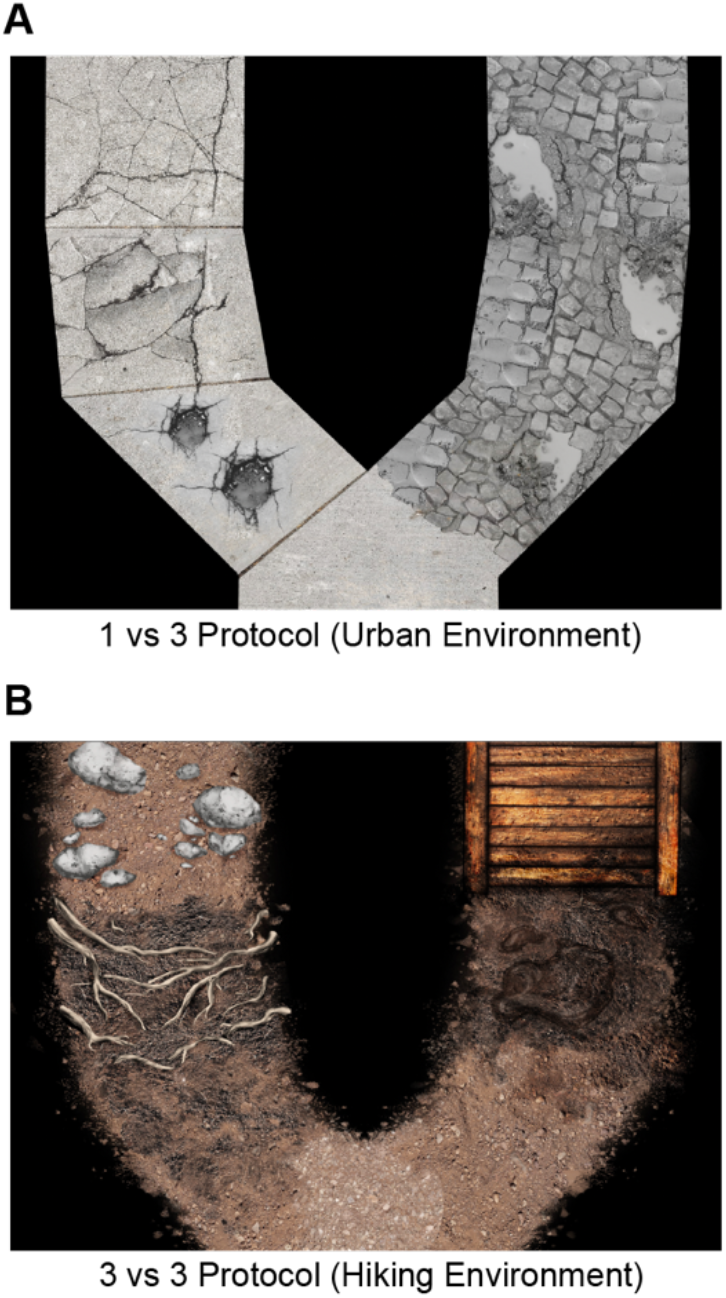
Examples of the environments used in this study. (A) An example of one of four urban environments used in the 1vs3 protocol. Terrain types include potholes (bottom left path), uneven sidewalk (middle left path), cracked sidewalk (top left path), and cobblestones with puddles (right path). None of the four hiking environments are shown. (B) An example of one of two hiking environments used in the 3vs3 protocol. Terrain types include damp dirt (bottom left path), tree roots (middle left path), rocks (top left path), dry dirt (bottom right path), mud (middle right path), and wooden bridge (top right path). None of the two urban environments are shown. See Fig. S1 in supplemental information for which terrain are used in the other environments in each protocol.

### Gaze behaviour is related to path choice

To show that gaze behaviour is involved in decision-making in our task, we compared the number of fixations and gaze time on the chosen and non-chosen paths. For the 1vs3 protocol, participants had a greater number of fixations on the chosen (7.8 ± 1.1) versus the non-chosen path (3.9 ± 0.9) (paired t-test: t_14_ = -10.6, p = 4.77e-8, Cohen’s d = 2.72). Participants also showed longer gaze time on the chosen (0.48 ± 0.05) versus the non-chosen path (0.20 ± 0.05) (paired t-test: t_14_ = - 13.6, p = 1.91e-9, Cohen’s d = 3.50). We found similar results for the 3vs3 protocol, where participants had a greater number of fixations on the chosen (7.5 ± 1.2) versus the non-chosen path (4.1 ± 1.0) (paired t-test: t_15_ = -9.8, p = 6.59e-8, Cohen’s d = 2.45), and longer gaze time on the chosen (0.45 ± 0.07) versus the non-chosen path (0.22 ± 0.05) (paired t-test: t_15_ = -8.4, p = 4.96e-7, Cohen’s d = 2.09). These results suggest that gaze is related to path choice.

### Gaze behaviour is related to information when looking at individual terrains, but not whole paths

We manipulated the number and type of terrain in each path to alter the amount of visual information present using two different approaches (1vs3 protocol and 3vs3 protocol); this allowed us to test whether a simple information-seeking strategy explains gaze decisions or self-confidence modifies information-seeking gaze behaviour. We first asked whether gaze is directed to the terrain images with the greatest amount of visual information, regardless of the path chosen. To address this question, we used the 3vs3 protocol, where we determined marginal (or Shannon) entropy (ME) and mean information gain (MIG) for each patch of terrain in each environment based on its greyscale image, the three components of the CIE-L*a*b* colour space, and the three components—hue, saturation, value—of the HSV colour space (Proulx and Parrott 2008). Greater amount of visual information contained within the terrain images equates to greater values of MIG and ME. Tables S1 and S2 in supplemental information show MIG and ME values, respectively, for each terrain of each environment. Examples of different terrains and their respective MIG_hue_ values are found in Fig. 2A. We determined the relationship between these information metrics and number of fixations and gaze time on each terrain. We found that increases in the MIG of the hue component (MIG_hue_) of a terrain significantly associated with increases in number of fixations (R^2^ = 0.22, ß = 4.82 [95% CI: 3.89, 5.75], p = 1.40e-21) and gaze time (Fig. 2B; R^2^ = 0.19, ß = 1.17 [95% CI: 0.93, 1.41], p = 9.89e-20). Greater ME of the hue component (ME_hue_) of a terrain also significantly associated with increases in number of fixations (R^2^ = 0.13, ß = 1.79 [95% CI: 1.30, 2.28], p = 2.86e-12), and gaze time (R^2^ = 0.09, ß = 0.40 [95% CI: 0.28, 0.53], p = 8.79e-10). However, the statistical model for MIG_hue_ had a lower AIC value than the model with ME_hue_ for number of fixations (ΔAIC = 43.7) and gaze time (ΔAIC = 45.3). Consequently, we used MIG_hue_ for subsequent analyses. No other colour-space information metric predicted gaze behaviour (see Tables S3 and S4 in supplemental information). Thus, there is only limited evidence to indicate that gaze is attracted to greater visual information regions within the paths.

**Figure 2:**
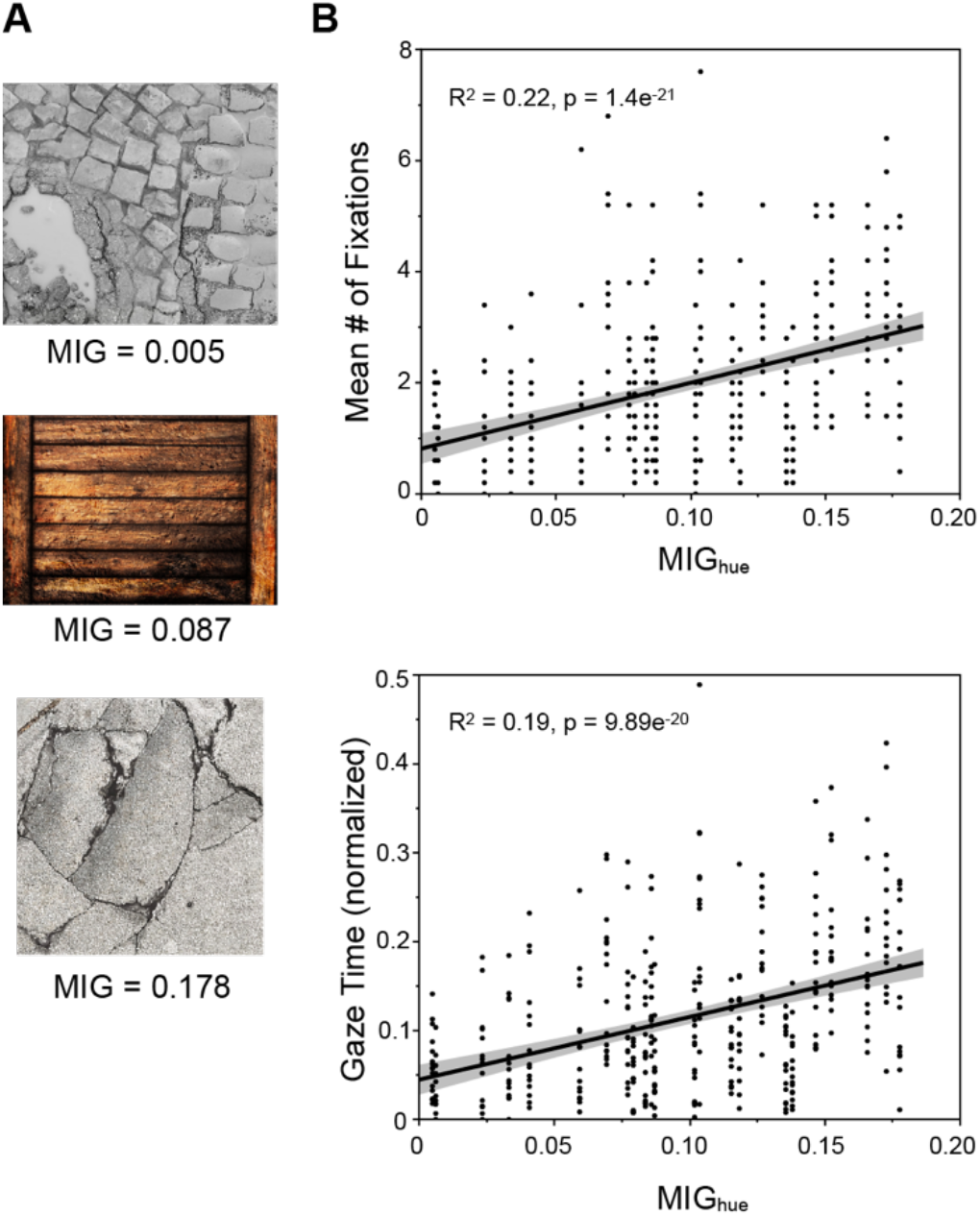
Relationship between mean information gain (MIG) and gaze behaviour during the approach (decision-making) phase in the 3vs3 protocol. (A) Example terrain image with low (top), medium (middle), and high (bottom) MIG based on hue. (B) Scatterplots of the number of fixations (top) or gaze time (bottom) and MIG based on hue in a terrain image. Gaze time is normalized to approach phase duration. In each scatterplot, solid black lines show the linear fits obtained from the models and grey shaded regions represent the 95% confidence intervals.

We next asked whether the amount of visual information across the entire path influences gaze. For the 1vs3 protocol, we defined information based on the number of terrains in each path. Specifically, we assumed that the path with three types of terrain (non-uniform) contained more visual information than the path with a single type of terrain (uniform). From a simple information-seeking perspective, we would predict that gaze is directed more frequently (and for longer duration) to the non-uniform path. We ran separate paired t-tests comparing the number of fixations and gaze time on each type of path (Fig. 3A). Participants made a similar number of fixations on both paths (uniform path = 5.84 ± 0.80; non-uniform path = 5.90 ± 0.97; paired t-test: t_14_ = -0.21, p = 0.835, Cohen’s d = 0.05). Participants also had similar gaze times on each path (uniform path = 0.34 ± 0.06; non-uniform path = 0.34 ± 0.04; paired t-test: t_14_ = 0.35, p = 0.733, Cohen’s d = 0.09). Thus, the amount of information did not attract gaze in this protocol.

**Figure 3:**
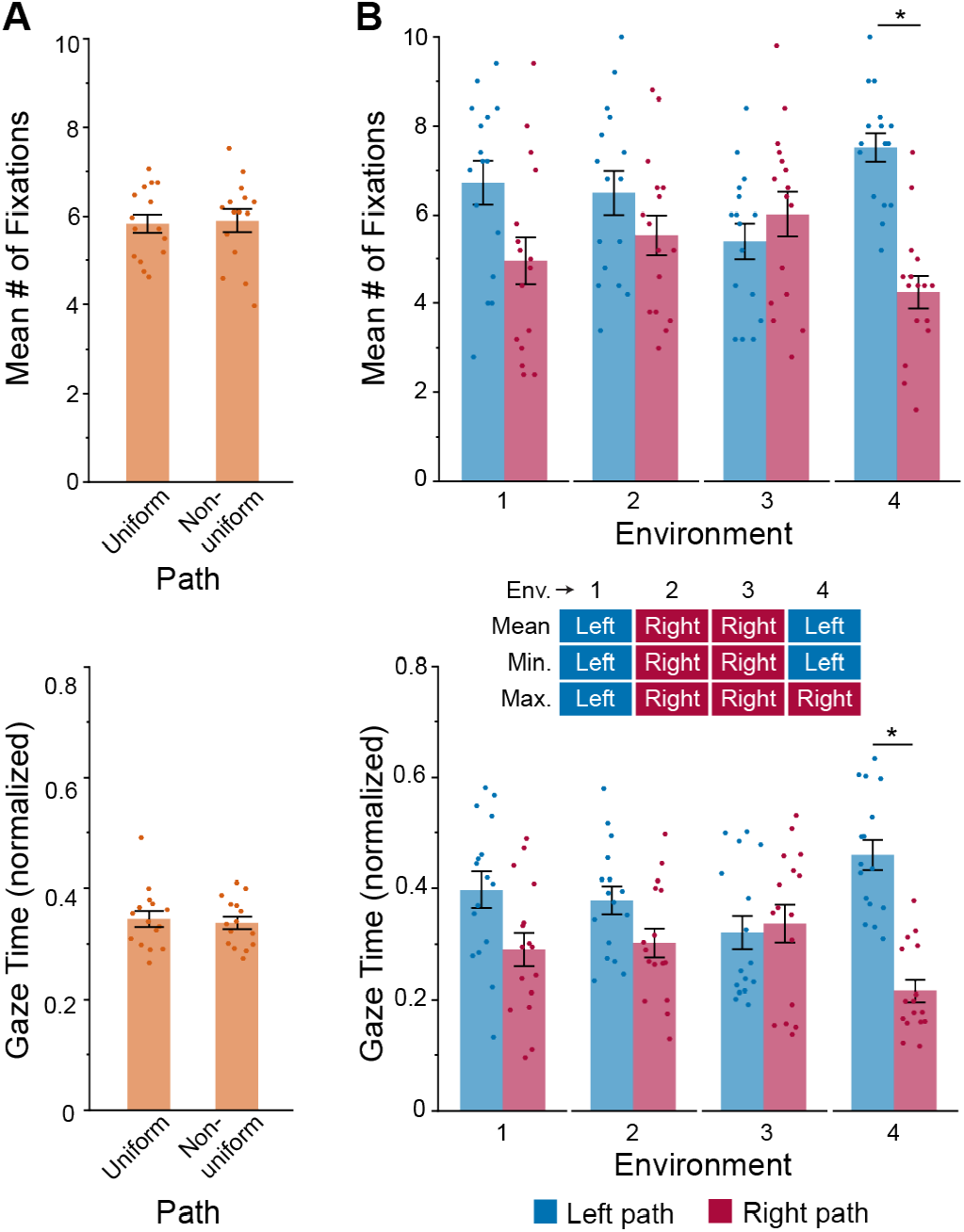
How the amount of information (number of terrain types or MIGhue) within a path choice relates to gaze behaviour during the approach (decision-making) phase. (A) Group mean ± SE number of fixations (top) or gaze time (bottom) on the path with low information (1 terrain type; uniform path) and the path with high information (3 terrain types; non-uniform path). (B) Group mean ± SE number of fixations (top) or gaze time (bottom) on the left and right path options. Inset: the chart shows which path, in each environment (env.), had greater MIGhue based on the mean of each terrain, the minimum (min.) terrain, or the maximum (max.) terrain. Gaze time is normalized to approach phase duration. Mean individual participant (n = 15 or 16, depending on the protocol) data values are superimposed. Asterisk indicates a statistically significant difference between the left and right path (p < 0.05).

For the 3vs3 protocol, if gaze is biased to the path with greater information, we would expect a greater frequency (and longer duration) to the left or right path depending on which one contained more visual information (i.e., greater MIG). For a given path, a person may rely on an average (or sum of) information across the three terrain types, the most informative terrain, or the least informative terrain to make decisions. Because we did not know a priori which method the brain uses, we used the MIG_hue_ for each terrain and determined for each environment which path had the most amount of information based on these three approaches. The inset chart in Figure 3B summarizes which path contained the most information for each environment. Except for environment 4 (Fig. 3B), we did not detect significant differences between the left and the right path in the number of fixations (paired t tests: environment 1, t_15_ = -1.89, p = 0.078; environment 2, t_15_ = -1.17, p = 0.260; environment 3, t_15_ = 0.75, p = 0.467) or gaze time (paired t tests: environment 1, t_15_ = -1.80, p = 0.092; environment 2, t_15_ = -1.61, p = 0.129; environment 3, t_15_ = 0.25, p = 0.802). For environment 4, participants fixated the left path more frequently (paired t test: t_15_ = -5.85, p = 3.17e-5) and for longer duration (paired t test: -5.65, p = 4.63e-5) than the right path. Overall, our results demonstrate that visual information, by itself, does not account for the observed gaze behaviour.

### Self-confidence about walking across the terrain influences gaze behaviour

Gaze is directed more to the chosen path. Here we asked whether participants select the path based on self-confidence. We averaged terrain self-confidence scores on each path for every participant to determine which path in each environment they felt most self-confident about. Participants chose the path they had the most confidence on 71% of the time for the 1vs3 protocol (*χ*^2^: p = 2.7e-5) and 73% of the time for the 3vs3 protocol (*χ*^2^: p = 4.6e-6). Individual self-confidence scores are shown in Table S5 in supplemental information.

Given that gaze is directed more to the chosen path, and participants select the path they are more self-confident about walking across more frequently, we tested the hypothesis that self-confidence about the path affects where and for how long gaze is directed. For the 1vs3 protocol, if confidence about the terrain affects gaze, we expected participants to vary the frequency and duration of gaze on the non-uniform path based on their self-confidence about walking across the uniform path. Thus, we determined the relationship between the confidence level on the uniform path and number of fixations and gaze time on the non-uniform path. Self-confidence on the uniform path predicted the number of fixations made on the non-uniform path (R^2^ = 0.31, ß = -0.26 [95% CI: - 0.43, -0.10], p = 0.002), but not gaze time (R^2^ = 0.03, ß = -0.008 [95% CI: -0.019, 0.002], p = 0.129). Specifically, participants made a greater number of fixations on the non-uniform path as self-confidence on the uniform path decreased (Fig. 4A).

**Figure 4:**
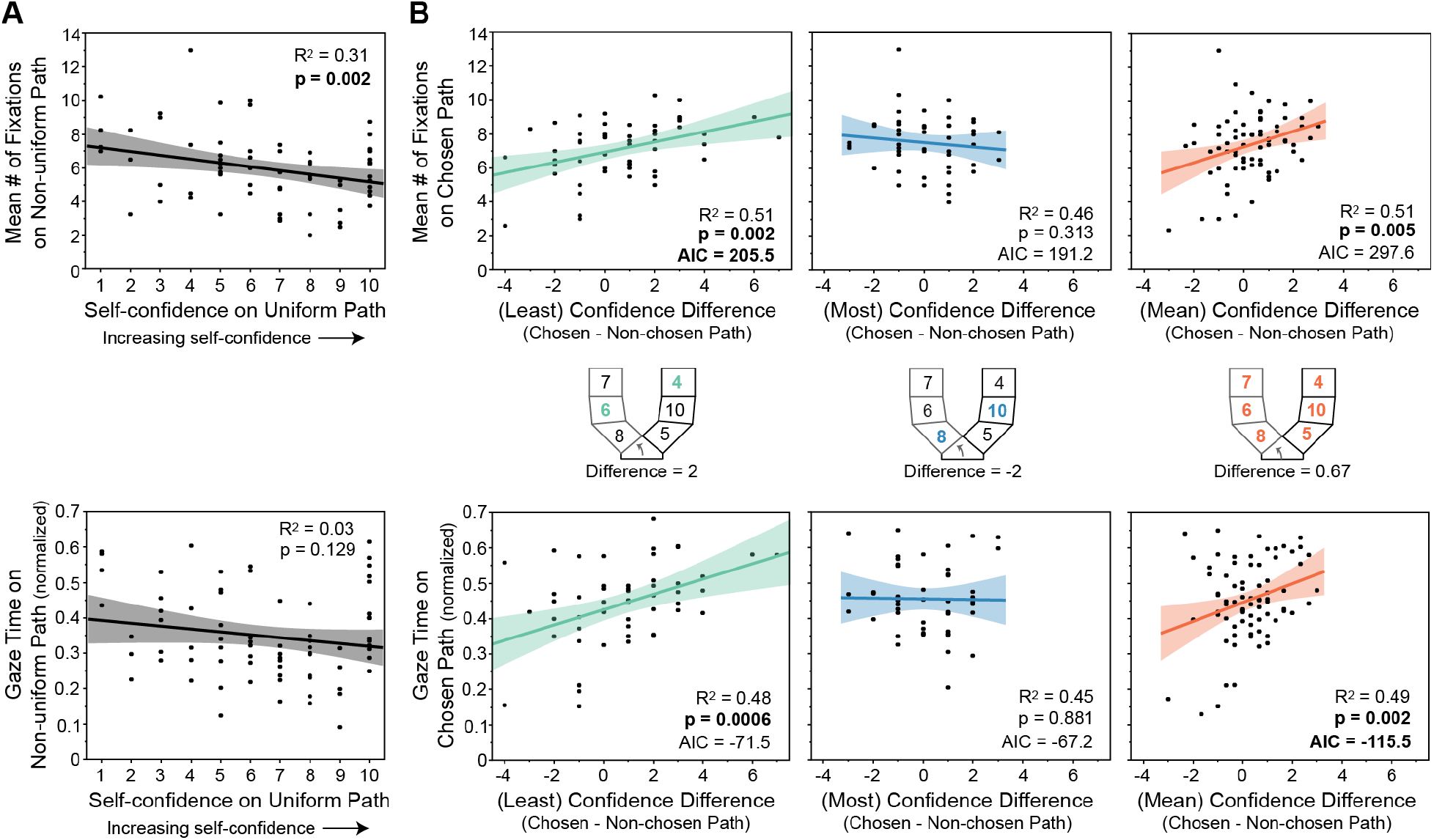
How self-confidence about walking across a path influences gaze behaviour during the approach (decision-making) phase. (A) Scatterplots of the number of fixations (top) or gaze time (bottom) on the non-uniform path versus the self-confidence level on the uniform path. (B) Scatterplots of the number of fixations (top) or gaze time (bottom) on the chosen path versus the difference in confidence between the two paths. Inset: illustrations of how confidence difference is calculated. Each section represents a terrain patch, with a confidence score inside (larger scores equal greater self-confidence in walking across it). The arrows point toward the chosen path (which has grey sections as opposed to black outlined sections). The coloured terrain numbers indicate which terrain is used in the calculation, which is based on the least confident terrain of each path (left), the most confident terrain of each path (middle), or the mean confidence across all terrains within a path (right). Gaze time is normalized to approach phase duration. In each scatterplot, solid lines show the linear fits obtained from the models and shaded regions represent the 95% confidence intervals.

For the 3vs3 protocol, both paths consisted of three different types of terrain. For each environment, we calculated the difference in self-confidence ratings between the chosen and non-chosen paths to determine whether people consider the difference in how self-confident they are about the terrain in each path when making gaze decisions. We used this metric because the brain can assign and compare values for each choice when making decisions (Rangel et al. 2008), and changes in gaze behaviour (i.e., saccade vigor) occur as a function of the difference in the subjective value that a participant assigns to each of two options (Reppert et al. 2015). Like information, a person may calculate confidence level on a given path based on an average (or sum of) confidence across the three terrain types, the most confident terrain of each path, or the least confident terrain of each path. Again, because we did not know a priori which method the brain uses, we tested models of each. The difference in the least confident terrain between paths predicted gaze behaviour (left column of Figure 4B; number of fixations: R^2^ = 0.51, ß = 0.29 [95% CI: 0.11, 0.46], p = 0.002; gaze time: R^2^ = 0.48, ß = 0.023 [95% CI: 0.010, 0.035], p = 0.0006).

The difference in the mean confidence across the terrain in each path also predicted gaze behaviour (right column of Figure 4B; number of fixations: R^2^ = 0.51, ß = 0.41 [95% CI: 0.13, 0.70], p = 0.005; gaze time: R^2^ = 0.49, ß = 0.029 [95% CI: 0.011, 0.046], p = 0.002). For the number of fixations, the linear mixed effects model using the difference in least confident terrain better explained the data (ΔAIC = 92.1). For gaze time, however, the linear mixed effects model using the difference in mean confidence better explained the data (ΔAIC = 44). In both cases, a greater difference in confidence (i.e., greater confidence with the eventually chosen path compared to the non-chosen path) associated with more fixations and longer gaze times on the chosen path. Taken together, our results demonstrate that the brain considers self-confidence when deciding where to direct gaze.

## DISCUSSION

The brain must decide which sources of information to sample via gaze to make the best possible decision about an action. Previous work shows that confidence in making the correct choice in perceptual decision or multi-arm bandit tasks influences information seeking-behaviour (Boldt et al. 2019; Desender et al. 2018, 2019; Pescetelli et al. 2021). However, when dealing with decisions related to motor behaviour, one’s self-confidence in performing the action of choice is arguably a more appropriate version of confidence. Here we tested the hypothesis that self-confidence about walking across a path affects where and for how long gaze is directed to each path in addition to the choice of which path to take. We used a forced-choice paradigm, where we manipulated the types of terrain in each path so that participants would have different levels of self-confidence about walking on them. Participants looked more to the path they eventually chose to walk across, which was usually the path they had greater self-confidence about. Although MIG of hue in the terrain predicted the frequency and duration of gaze on a given terrain patch, participants did not direct gaze more to the overall path with greater information. Rather, we found that less confidence on the uniform (single) terrain path associated with a greater number of fixations on the non-uniform (three) terrain path. With two non-uniform paths, greater self-confidence with the path eventually chosen, relative to the alternative, associated with a greater number of fixations and gaze time on it during the approach. Taken together, our results demonstrate that the brain uses self-confidence to guide gaze and walking decisions in complex environments.

A potential limitation of our study is that we projected images of terrain on the ground rather than having participants walk across real terrain. We chose this method for two reasons. One, our lab-based, simulated terrain provided us with greater control over the design of the walking paths. Two, the projected images eliminate (or reduce) any energetic and/or stability cost associated with walking across real terrain. This latter point is important since both motor costs influence gaze behaviour (Domínguez-Zamora and Marigold 2019, 2021; Matthis et al. 2018; Moskowitz et al. 2023), and we wanted to demonstrate a role for self-confidence. We believe our results generalize to the real world because despite the simulated terrain, participants usually chose the path they had more confidence in walking across (>70% of trials) and often avoided stepping on the images of rocks, potholes, puddles, and mud when choosing where to place their feet on the paths.

The amount of available visual information contained within a patch of terrain, but not the walking path, predicted gaze allocation. In natural behaviour, the brain appears to direct gaze to gain information and solve task-relevant uncertainties about the environment (Hayhoe 2017; Sprague and Ballard 2003; Sprague et al. 2007). Indeed, Domínguez-Zamora et al. (2018) found that having to step to the centre of targets led to longer gaze times when the target was blurred (and thus the centre location more uncertain). Furthermore, Tong et al. (2017) found a greater number of fixations to task-relevant obstacles to avoid when their random motion increased the uncertainty of their position. Thus, we reasoned that one possibility in our study was that participants would direct gaze more frequently (or for longer duration) to individual patches of terrain (or the entire walking path) with greater amounts of visual information to reduce uncertainty associated with that location. We defined the amount of information as either the MIG (or ME) across different colour spaces of each terrain (3vs3 protocol) or the number of terrain patches along a path (1v3 protocol). At the level of individual terrain (3vs3 protocol), we found a significant association between MIG_hue_ and number of fixations and gaze time on a patch of terrain. Hue is a representative measure on how the human visual system perceives wavelengths in the light signal (Proulx and Parrott 2008). The brain may initially perceive each object/surface in the environment based on their dominant hue (Neitz & Neitz 2008; Stoughton and Conway 2008). By simplifying surfaces to their hue, the brain can faster differentiate between them (Borstein and Korda 1984), aiding in object recognition and calculations of the number of different surfaces in an environment (Valberg 2001). However, we found no clear relationship between gaze and our metrics of information (with either protocol) when considering the entire path. It is important to acknowledge that there are other ways to quantify visual information besides ME and MIG, which might show a relationship to gaze metrics. We opted to use MIG and ME as they are common techniques when determining information from images of natural environments (Proulx and Parrott 2008). Taken together, our results suggest that other factors beyond information mediate gaze decisions.

Self-confidence about walking on specific terrain influenced how people directed gaze to gain information. Specifically, when there was an equal number of different terrains within each of two path options, we showed that people considered the difference in their self-confidence between paths (the difference between the least confident terrains or between mean confidence across the terrains). The greater the self-confidence in the path they eventually chose relative to the non-chosen path, the more people directed gaze to the chosen path. When one path option had a greater number of terrain types, we demonstrated that people were more likely to visual explore this path when they were less self-confident with the alternative. If self-confidence about an option is high, there is less need to seek out additional information to inform the decision of which path to take. In this sense, people will exploit (i.e., direct gaze more to) the path option they have greater self-confidence about walking across. These results are in line with previous research that operationally defined confidence as the certainty of a correct response (Desender et al. 2018; Pescetelli et al. 2021). In these perceptual decision-making paradigms, participants were more likely to choose to view a simpler version of a stimulus again (Desender et al. 2018) or seek advice about the correct answer (Pescetelli et al. 2021) when they were less confident about their initial judgement. Thus, self-confidence about one’s ability to perform a motor action and confidence in a decision appear to influence behaviour in a similar manner and as a result, may share similar neural substrates.

How do our results relate to models of gaze and decision making? Like with a choice between two foot-placement targets (Domínguez-Zamora and Marigold 2021), we found that participants looked more to the chosen walking path. This is consistent with predictions of the attentional drift-diffusion and gaze-weighted linear accumulator models (Krajbich et al. 2010; Thomas et al. 2019). These models assume that momentary gaze directed to an item introduces a choice bias for that option, and when not looking at an item, its decision value is discounted. However, these models were not designed to address motor behaviour, where choices may change as movement unfolds, movements have consequences (which may affect balance), and where a person must decide how to act based on accumulated information from gaze. A more suitable model to compare our results with is one developed by Sprague and colleagues (Sprague and Ballard 2003; Sprague et al. 2007), which aims to explain gaze allocation during a motor behaviour that is divided into subtasks. The model assigns a value to a saccade based on the expected negative consequences of the uncertainty created when a fixation is not made to a specific task-relevant location. Gaze is directed to reduce the uncertainty (or gain information) about aspects of the environment where it serves to maximize a reward associated with an action or task goal. In our study, however, participants did not direct gaze more to the path with greater visual information and thus, would not have reduced task-relevant uncertainty as much as possible. Neither the evidence accumulation models described above, nor this reinforcement learning-based model address the fact that self-confidence can bias gaze. Our results suggest that decision-making models of gaze should incorporate the idea of self-confidence, at least when trying to explain motor behaviour.

Overall, our work contributes to the growing understanding of how different types of confidence affect different domains (e.g., perceptual, value-based) of decision-making (Boldt et al. 2019; Desender et al. 2018; De Martino et al. 2013; Kepecs and Mainen 2012). Here we show that self-confidence is one of the brain’s inputs in the decision-making process that guides gaze and walking behaviours. More specifically, self-confidence in one’s motor abilities predicts how information about the environment in gathered via gaze in addition to one’s choice of walking path to take. Decisions related to motor behaviour are ubiquitous. Since movement choices have consequences, such as whether the intended goal is achieved, or whether they are benign or can lead to potential harm, one’s self-confidence is likely to impact one’s decision about where or how to move. Thus, our work has broad implications about how we gather information and make decisions. Future work should determine additional factors that can modify information-seeking gaze behaviour and how they interact with self-confidence.

## ACKNOWLEDGEMENTS

The authors thank Ian Bercovitz for advice on statistical analyses and Laura Gimenes for help with terrain illustrations.

## METHODS

### Participants

Sixteen healthy adults participated in this study (5 women and 11 men; mean age = 26 ± 3 years). Due to problems with recording gaze, we excluded one participant for one block of trials. Participants did not have any known visual, neurological, muscular, or joint disorder that could affect their walking or gaze behaviour. The Office of Research Ethics at Simon Fraser University approved the study protocol, and participants provided informed written consent prior to participating.

### Experimental design

Participants performed a visually guided walking paradigm that required them to walk across one of two paths that we projected on the ground. Using images projected on the ground ensured that energetic- and stability-related motor costs did not confound our results, as both can influence gaze and walking decisions (Domínguez-Zamora and Marigold 2019, 2021; Matthis et al. 2018). We created the path images with Photoshop (Adobe Inc., San Jose, CA, USA). Each path was approximately 2 meters long and 55 cm wide and joined at the start (see Fig. 1). We used two different environmental themes to create our paths: hiking and urban. For the hiking environment, we used terrain patches commonly found on hiking trails, such as mud, tree roots, dirt, and rocks. For the urban environment, we used asphalt, cobblestones, and concrete. Paths in the same projection always shared a common theme, so that participants never faced a scenario where they had one hiking path and an urban path in the same environment.

We configured the environments in MATLAB (The MathWorks, Natick, MA) with the Psychophysics Toolbox, version 3 (Brainard 1997). An LCD projector (Epson PowerLight 5535U, brightness of 5,500 lumens) displayed the environments on a black uniform mat. To diminish the effect of environmental references and increase image visibility, participants walked under reduced light conditions (range of 1.1 to 3.5 lux, like a moonlit night). We recorded kinematic data at 100 Hz using two Optotrak Certus motion capture cameras (Northern Digital, Waterloo, ON, Canada) positioned perpendicular to the walking path. This involved recording infrared-emitting position markers placed on the participant’s head and chest and bilaterally on each midfoot (second to third metatarsal head). We also recorded gaze data at 100 Hz using a high-speed mobile eye tracker (Tobii Pro Glasses 3, Tobii Technology Inc., Reston, VA, USA) mounted on the participant’s head and synchronized with the motion capture system. We calibrated the eye tracker before each of the two protocols.

### Experimental protocol

Before the start of the experimental trials, we presented participants with images on the ground of the 12 types of terrains we used to construct the environments. This allowed the participants to become familiar with the different terrain. Subsequently, we had participants complete two different protocols (see Fig. 1), the order of which was counterbalanced. In one protocol, referred to as 1vs3, we used eight different environments (four hiking and four urban), and participants completed 32 walking trials (four trials of each environment in random order). Environments had two different paths of the same setting (hiking or urban), where one path consisted of one type of terrain (uniform path) and one path consisted of three types of terrain (non-uniform path). In the other protocol, referred to as 3vs3, we used four different environments (two hiking and two urban), and participants completed 20 walking trials (five trials of each environment in random order). Environments had two different paths of the same setting (hiking or urban), which each consisted of three types of terrain. See Fig. S1 in supplemental information for which terrain are used in the environments in each protocol.

In both protocols, for each walking trial, participants started from a standing position approximately 1.5 m from the projected walking paths. We projected a fixation cross at the centre of the projection area, approximately 2 meters from the starting point and instructed participants to maintain their gaze on it until the image of the environment appeared. After one second, the cross disappeared and one of the environments appeared. We asked participants to remain stationary and freely visually explore the terrains for two seconds. In real life, a person would likely see their path choices well in advance and could visually explore as they approach from a distance. Our choice to allow participants two seconds to explore from a stationary position substituted for this ability, since we did not have the walking space for participants to complete more than 2 steps before reaching one of the paths. After the two seconds, an auditory cue signalled participants to begin walking. We asked participants to pretend the terrains were real, to choose the path they would normally take as if outside of the lab, and to step where they would normally step if they faced that terrain in real life. We instructed participants to walk at a self-selected speed and stop two steps after walking across their chosen path. An experimenter recorded which path the participant chose on each trial during both protocols.

After the experimental walking trials, we had each participant rate their confidence level of each terrain. Specifically, we asked participants: “for each type of terrain, please indicate how confident (or certain) you are of walking across it without losing balance, as though you had to step on it in real life outside. Please use a scale of 1 to 10 (where 1 is not at all confident and 10 is extremely confident).”

### Data and statistical analyses

We filtered kinematic data using a 6-Hz low-pass Butterworth algorithm. We used this data to calculate the approach phase (defined as the time between the start of the trial and the participant’s first foot contact with the fork in the joined paths). Because we were interested in understanding the decision-making process, we only analyzed gaze behaviour during the approach phase.

To analyze gaze data, we used GlassesViewer (Niehorster et al. 2020). We defined fixations as the times during which a target or region on the ground stabilized on the retina and detected them based on the slow-phase classifier described in Hessels et al. (2020). This classifier uses an adaptive velocity threshold based on estimated gaze velocity. We used the following classifier parameters: 5000 deg/s start velocity threshold; 50 ms minimum fixation duration; lambda slow/fast separation threshold of 2.5; 8 s moving window. We used the 30 Hz eye tracker video with the gaze location superimposed on the image to verify the presence and location of fixations. To quantify gaze behaviour, we calculated the number of fixations and gaze time (i.e., sum of fixation times) on each terrain. We normalized gaze time on each terrain by the approach time to control for any differences in gait initiation and speed across trials and participants.

For both protocols, we first sought to demonstrate a link between gaze behaviour and path choice (decision-making). Specifically, we compared the number of fixations (or gaze time) during the approach phase on the chosen path with the non-chosen path using separate paired t tests. We predicted a greater number of fixations and gaze times on the chosen path.

We manipulated the number and type of terrain in each path to alter the amount of visual information present using two different approaches (1vs3 protocol and 3vs3 protocol). For the 1vs3 protocol, we assumed that the non-uniform path (with three types of terrain) contains more visual information than the uniform path (with one type of terrain). Thus, we compared the number of fixations (or gaze time) on the uniform and non-uniform paths using paired t tests. Next, we tested whether the level of confidence associated with a path predicted gaze behaviour on a path. Specifically, we performed a linear mixed-effects model, with the number of fixations on the non-uniform path as the response variable and level of confidence on the uniform path as the predictor variable. We included participant as a random effect. We also performed a linear regression, with gaze time on the non-uniform path as the response variable and level of confidence on the uniform path as the predictor variable. In this model, we removed the random effect of participant because the variance component of this effect was not above zero. For each model, we included each participant’s perceived self-confidence, thus accounting for individual perceptions. We used confidence on the uniform path, rather than the non-uniform path, because only one type of terrain is present and is thus easier to interpret. We used a chi-square test to determine if participants chose the more confident path more frequently.

For the 3vs3 protocol, we quantified information as the information contained in the visual image of each patch of terrain using several different metrics (Proulx and Parrott 2008). Specifically, we calculated marginal (or Shannon) entropy (ME) as:

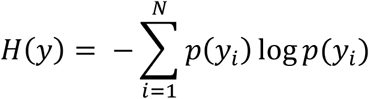

where *p(y*_*i*_) is the probability of observing a pixel value independent of its position in the image and *N* is the number of bins of pixel values. In addition, we calculated the mean information gain (MIG), as described in detail in Proulx and Parrott (2008) and based on Andrienko et al. (2000). MIG requires calculation of the joint entropy, which is defined as:

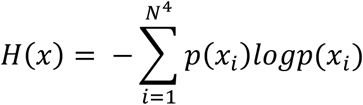

where *p(x*_*i*_) is the probability of finding a 2 x 2 colour combination x_i_ in the image and *N*^*4*^ is the number of theoretical combinations. MIG is the joint entropy minus the marginal entropy and specified as:

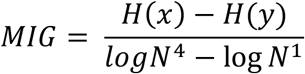

where we used an *N* = 8. MIG is zero for uniform (ordered) patterns and is 1 for random (disordered) patterns. High values of MIG represent high information gain. Due to the curvature of each path, we extracted the maximum rectangular area of each terrain patch for each environment to run the information analysis on. This yielded 24 terrain patches (6 terrains/environment x 4 environments).

We determined ME and MIG for each patch of terrain in each environment based on its greyscale image, the three components of the CIE-L*a*b* colour space, and the three components—hue, saturation, value—of the HSV colour space using the *MIG_simple*.*m* routine from Proulx and Parrott (21). The CIE-L*a*b* colour space is based on the human visual system and the opponent process theory of colour vision, while the HSV colour space is based on how humans perceive colour. Next, we used a series of linear regressions or linear mixed-effects models, with either the mean number of fixations or the gaze time (with a square root transformation to ensure normality) on the patch of terrain as the response variable, one of the information metrics as the predictor variable, and participant as a random effect if the variance component of this effect was > 0. Due to the number of tests with this analysis, we used a more conservative alpha level of 0.01, as opposed to 0.05 for all other statistical analyses. We found two statistically significant models, for ME and MIG associated with hue (see Results), but only used the latter in our subsequent analysis because the model had a lower Akaike information criterion (AIC). We asked whether gaze is directed more to the path with greater information. Specifically, we determined whether the left or right path of each of the four environments had greater information based on the average (or sum of) information across the three terrain types, the most informative terrain, and the least informative terrain. Subsequently, we performed separate paired t-tests comparing mean number of fixations or gaze time on the left and right paths for each environment.

For this (3vs3) protocol, for each environment, we calculated the difference in self-confidence ratings between the two paths. Like information, a person may calculate confidence level on a given path based on an average (or sum of) confidence across the three terrain types, the most confident terrain, or the least confident terrain. Again, because we did not know a priori which method the brain uses, we tested models of each and compared statistically significant ones using the AIC. Specifically, we used linear mixed-effects models, with the number of fixations (or gaze time) on the chosen path as the response variable and the difference in self-confidence ratings between paths as the predictor variable. In all cases, we included participant as a random effect. We used chi-square tests to determine if participants chose the more confident path (based on the average, or sum of, confidence across the three terrain types, the most confident terrain, or the least confident terrain) more frequently.

