## Supplemental Information for "Fork in the road: how self-confidence about terrain influences gaze behaviour and path choice"

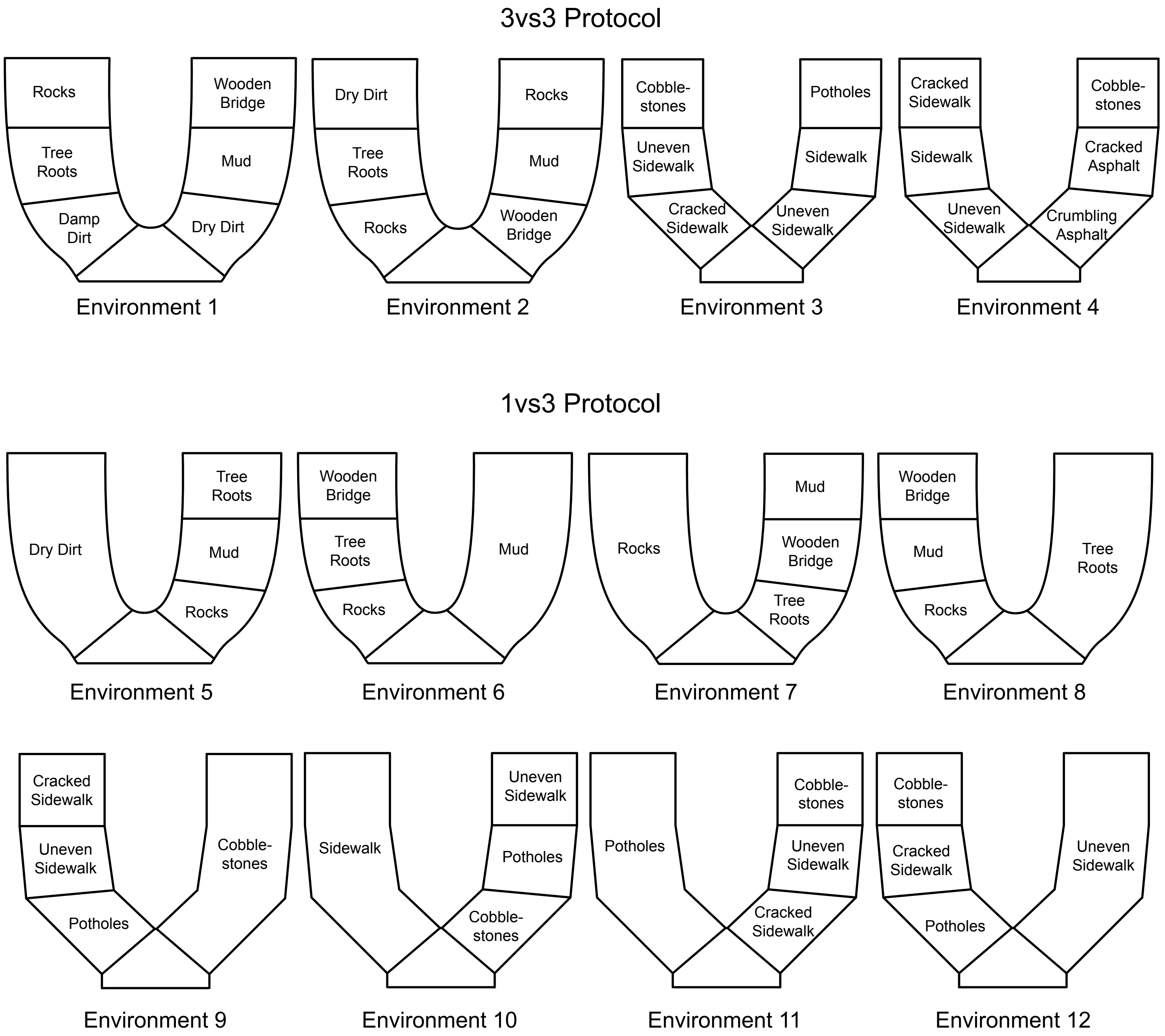


**Figure S1:** Terrain layout for each environment for the 3vs3 and 1vs3 protocols. See also Figure 1.

**Table S1:** Mean information gain (MIG) for each terrain and environment.

| Terrain Code | $MIG_{grey}$ | $MIG_{L*}$ | $MIG_{a*}$ | $MIG_{b*}$ | $MIG_{hue}$ | $MIG_{saturation}$ | $MIG_{value}$ |
|---|---|---|---|---|---|---|---|
| env1p1 | 0.53 | 0.55 | 0.31 | 0.28 | 0.06 | 0.46 | 0.55 |
| env1p2 | 0.55 | 0.56 | 0.27 | 0.25 | 0.17 | 0.54 | 0.57 |
| env1p3 | 0.47 | 0.46 | 0.21 | 0.18 | 0.04 | 0.31 | 0.46 |
| env1p4 | 0.55 | 0.55 | 0.30 | 0.26 | 0.02 | 0.46 | 0.56 |
| env1p5 | 0.44 | 0.46 | 0.28 | 0.27 | 0.15 | 0.62 | 0.47 |
| env1p6 | 0.50 | 0.51 | 0.35 | 0.33 | 0.09 | 0.65 | 0.53 |
| env2p1 | 0.42 | 0.43 | 0.22 | 0.19 | 0.07 | 0.29 | 0.43 |
| env2p2 | 0.55 | 0.56 | 0.23 | 0.26 | 0.15 | 0.53 | 0.57 |
| env2p3 | 0.52 | 0.52 | 0.30 | 0.26 | 0.01 | 0.42 | 0.52 |
| env2p4 | 0.45 | 0.46 | 0.34 | 0.33 | 0.08 | 0.58 | 0.49 |
| env2p5 | 0.43 | 0.45 | 0.31 | 0.29 | 0.17 | 0.63 | 0.46 |
| env2p6 | 0.50 | 0.51 | 0.22 | 0.20 | 0.10 | 0.39 | 0.51 |
| env3p1 | 0.58 | 0.58 | 0.28 | 0.25 | 0.14 | 0.03 | 0.58 |
| env3p2 | 0.54 | 0.54 | 0.19 | 0.10 | 0.13 | 0.03 | 0.54 |
| env3p3 | 0.34 | 0.32 | 0.05 | 0.05 | 0.03 | 0.01 | 0.34 |
| env3p4 | 0.57 | 0.55 | 0.20 | 0.19 | 0.18 | 0.05 | 0.56 |
| env3p5 | 0.35 | 0.32 | 0.17 | 0.07 | 0.08 | 0.00 | 0.35 |
| env3p6 | 0.37 | 0.36 | 0.09 | 0.07 | 0.08 | 0.09 | 0.38 |
| env4p1 | 0.60 | 0.59 | 0.21 | 0.25 | 0.10 | 0.06 | 0.60 |
| env4p2 | 0.34 | 0.31 | 0.17 | 0.14 | 0.09 | 0.01 | 0.34 |
| env4p3 | 0.69 | 0.67 | 0.24 | 0.22 | 0.12 | 0.08 | 0.68 |
| env4p4 | 0.61 | 0.62 | 0.10 | 0.14 | 0.14 | 0.03 | 0.61 |
| env4p5 | 0.63 | 0.65 | 0.12 | 0.12 | 0.12 | 0.09 | 0.63 |
| env4p6 | 0.34 | 0.35 | 0.005 | 0.01 | 0.005 | 0.0002 | 0.34 |

Env = environment (1 to 4); p = position along path (p1 = proximal terrain on left path; p2 = middle terrain on left path; p3 = distal terrain on left path; p4 = proximal terrain on right path; p5 = middle terrain on right path; p6 = distal terrain on right path); grey = greyscale; L* = perceptual lightness of CIE L*a*b* colour space; a* = red/green scale of CIE L*a*b* colour space; b* = blue/yellow scale of CIE L*a*b* colour space; hue = hue component of HSV colour space; saturation = saturation component of HSV colour space; value = value component of HSV colour space.

**Table S2:** Marginal entropy (ME) for each terrain and environment.

| Terrain Code | $ME_{grey}$ | $ME_{L*}$ | $ME_{a*}$ | $ME_{b*}$ | $ME_{hue}$ | $ME_{saturation}$ | $ME_{value}$ |
|---|---|---|---|---|---|---|---|
| env1p1 | 0.83 | 0.85 | 0.56 | 0.62 | 0.10 | 0.63 | 0.88 |
| env1p2 | 0.90 | 0.91 | 0.47 | 0.51 | 0.27 | 0.71 | 0.94 |
| env1p3 | 0.88 | 0.89 | 0.63 | 0.63 | 0.11 | 0.72 | 0.84 |
| env1p4 | 0.83 | 0.84 | 0.48 | 0.48 | 0.03 | 0.65 | 0.86 |
| env1p5 | 0.68 | 0.71 | 0.48 | 0.51 | 0.21 | 0.76 | 0.76 |
| env1p6 | 0.87 | 0.89 | 0.68 | 0.80 | 0.12 | 0.80 | 0.98 |
| env2p1 | 0.87 | 0.88 | 0.64 | 0.64 | 0.16 | 0.71 | 0.85 |
| env2p2 | 0.90 | 0.92 | 0.44 | 0.55 | 0.24 | 0.71 | 0.95 |
| env2p3 | 0.79 | 0.79 | 0.52 | 0.55 | 0.01 | 0.60 | 0.80 |
| env2p4 | 0.86 | 0.89 | 0.68 | 0.78 | 0.11 | 0.78 | 0.98 |
| env2p5 | 0.66 | 0.69 | 0.50 | 0.52 | 0.23 | 0.77 | 0.74 |
| env2p6 | 0.91 | 0.92 | 0.60 | 0.60 | 0.21 | 0.76 | 0.91 |
| env3p1 | 0.86 | 0.85 | 0.56 | 0.46 | 0.34 | 0.05 | 0.86 |
| env3p2 | 0.85 | 0.86 | 0.46 | 0.28 | 0.34 | 0.05 | 0.85 |
| env3p3 | 0.70 | 0.72 | 0.26 | 0.55 | 0.19 | 0.02 | 0.70 |
| env3p4 | 0.86 | 0.84 | 0.36 | 0.35 | 0.32 | 0.08 | 0.85 |
| env3p5 | 0.46 | 0.42 | 0.48 | 0.22 | 0.25 | 0.004 | 0.46 |
| env3p6 | 0.70 | 0.68 | 0.33 | 0.56 | 0.27 | 0.22 | 0.71 |
| env4p1 | 0.79 | 0.78 | 0.43 | 0.51 | 0.23 | 0.08 | 0.79 |
| env4p2 | 0.45 | 0.42 | 0.46 | 0.36 | 0.26 | 0.01 | 0.46 |
| env4p3 | 0.87 | 0.85 | 0.43 | 0.42 | 0.22 | 0.11 | 0.86 |
| env4p4 | 0.94 | 0.96 | 0.26 | 0.26 | 0.39 | 0.04 | 0.94 |
| env4p5 | 0.88 | 0.90 | 0.35 | 0.36 | 0.36 | 0.10 | 0.88 |
| env4p6 | 0.71 | 0.70 | 0.06 | 0.06 | 0.06 | 0.0003 | 0.71 |

Env = environment (1 to 4); p = position along path (p1 = proximal terrain on left path; p2 = middle terrain on left path; p3 = distal terrain on left path; p4 = proximal terrain on right path; p5 = middle terrain on right path; p6 = distal terrain on right path); grey = greyscale; L* = perceptual lightness of CIE L*a*b* colour space; a* = red/green scale of CIE L*a*b* colour space; b* = blue/yellow scale of CIE L*a*b* colour space; hue = hue component of HSV colour space; saturation = saturation component of HSV colour space; value = value component of HSV colour space.

**Table S3:** Relationship between the number of fixations (response variable) versus information metrics (predictor variables).

| Predictor | $R^2$ | Estimate [95% CI] | Intercept | p-value | AIC |
|---|---|---|---|---|---|
| $ME_{hue}$ | 0.13 | 1.79 [1.30,2.28] | 0.92 | 2.86e-12 | 558.6 |
| $MIG_{hue}$ | 0.22 | 4.82 [3.89,5.75] | 0.83 | 1.40e-21 | 514.9 |
| $ME_{saturation}$ | 7.8e-7 | -0.001 [-0.16,0.16] | 1.30 | 0.986 | 599.2 |
| $MIG_{saturation}$ | 4.7e-4 | 0.05 [-0.17,0.27] | 1.30 | 0.672 | 599.0 |
| $ME_{value}$ | 1.9e-3 | -0.17 [-0.56,0.22] | 1.43 | 0.398 | 598.5 |
| $MIG_{value}$ | 1.0e-3 | 0.17 [-0.37,0.72] | 1.21 | 0.529 | 598.8 |
| $ME_{L*}$ | 2.7e-3 | -0.20 [-0.58,0.18] | 1.45 | 0.310 | 598.2 |
| $MIG_{L*}$ | 3.4e-4 | 0.09 [-0.42,0.61] | 1.25 | 0.720 | 599.1 |
| $ME_{a*}$ | 2.4e-3 | -0.18 [-0.55,0.19] | 1.38 | 0.341 | 598.3 |
| $MIG_{a*}$ | 1.2e-3 | -0.20 [-0.79,0.38] | 1.34 | 0.500 | 598.8 |
| $ME_{b*}$ | 3.0e-3 | -0.17 [-0.48,0.14] | 1.34 | 0.284 | 598.1 |
| $MIG_{b*}$ | 2.7e-4 | 0.10 [-0.50,0.69] | 1.28 | 0.750 | 559.1 |
| $ME_{grey}$ | 3.2e-3 | -0.23 [-0.65,0.18] | 1.48 | 0.266 | 598.0 |
| $MIG_{grey}$ | 1.4e-4 | 0.06 [-0.48,0.60] | 1.26 | 0.818 | 559.2 |

AIC = Akaike information criterion; ME = marginal entropy; MIG = mean information gain; grey = greyscale; L* = perceptual lightness of CIE L*a*b* colour space; a* = red/green scale of CIE L*a*b* colour space; b* = blue/yellow scale of CIE L*a*b* colour space; hue = hue component of HSV colour space; saturation = saturation component of HSV colour space; value = value component of HSV colour space.

For the statistical analysis, data were square root transformed to ensure normality.

**Table S4:** Relationship between normalized gaze time (response variable) versus information metrics (predictor variables).

| Predictor | $R^2$ | Estimate [95% CI] | Intercept | p-value | AIC |
|---|---|---|---|---|---|
| $ME_{hue}$ | 0.09 | 0.40 [0.28,0.53] | 0.22 | 8.79e-10 | -494.6 |
| $MIG_{hue}$ | 0.19 | 1.17 [0.93,1.41] | 0.20 | 9.89e-20 | -539.9 |
| $ME_{saturation}$ | 1.5e-4 | 0.005 [-0.04,0.05] | 0.31 | 0.814 | -456.8 |
| $MIG_{saturation}$ | 1.1e-3 | 0.02 [-0.04,0.07] | 0.30 | 0.513 | -457.2 |
| $ME_{value}$ | 4.1e-4 | -0.02 [-0.12,0.08] | 0.32 | 0.692 | -456.9 |
| $MIG_{value}$ | 3.3e-3 | 0.08 [-0.06,0.22] | 0.27 | 0.260 | -458.0 |
| $ME_{L*}$ | 7.2e-4 | -0.03 [-0.12,0.07] | 0.33 | 0.600 | -457.0 |
| $MIG_{L*}$ | 1.9e-3 | 0.06 [-0.07,0.19] | 0.28 | 0.392 | -457.5 |
| $ME_{a*}$ | 1.7e-3 | -0.04 [-0.13,0.06] | 0.33 | 0.426 | -457.4 |
| $MIG_{a*}$ | 1.0e-4 | -0.01 [-0.16,0.13] | 0.31 | 0.846 | -456.8 |
| $ME_{b*}$ | 1.2e-3 | -0.03 [-0.11,0.05] | 0.32 | 0.490 | -457.2 |
| $MIG_{b*}$ | 1.7e-3 | 0.06 [-0.09,0.21] | 0.30 | 0.419 | -457.4 |
| $ME_{grey}$ | 9.5e-4 | -0.03 [-0.14,0.07] | 0.33 | 0.548 | -457.1 |
| $MIG_{grey}$ | 1.5e-3 | 0.05 [-0.08,0.19] | 0.28 | 0.453 | -457.3 |

AIC = Akaike information criterion; ME = marginal entropy; MIG = mean information gain; grey = greyscale; L* = perceptual lightness of CIE L*a*b* colour space; a* = red/green scale of CIE L*a*b* colour space; b* = blue/yellow scale of CIE L*a*b* colour space; hue = hue component of HSV colour space; saturation = saturation component of HSV colour space; value = value component of HSV colour space.

For the statistical analysis, data were square root transformed to ensure normality.

**Table S5:** Self-confidence scores.

| Participant | Wooden bridge | Tree Roots | Rocks | Mud | Dry Dirt | Damp Dirt | Sidewalk | Cracked Sidewalk | Cracked Asphalt | Crumbling Asphalt | Uneven Sidewalk | Cobblestones | Potholes |
|---|---|---|---|---|---|---|---|---|---|---|---|---|---|
| **Pc01** | 9 | 6 | 7 | 4 | 10 | 9 | 10 | 9 | 9 | 8 | 7 | 8 | 7 |
| **Pc02** | 8 | 4 | 4 | 2 | 9 | 8 | 10 | 9 | 9 | 6 | 6 | 3 | 4 |
| **Pc03** | 4 | 5 | 3 | 1 | 7 | 6 | 8 | 6 | 6 | 5 | 4 | 4 | 3 |
| **Pc04** | 10 | 5 | 8 | 3 | 9 | 7 | 10 | 10 | 10 | 9 | 8 | 7 | 5 |
| **Pc05** | 10 | 8 | 7 | 7 | 10 | 9 | 10 | 10 | 10 | 10 | 10 | 8 | 8 |
| **Pc06** | 10 | 8 | 8 | 6 | 10 | 9 | 10 | 10 | 10 | 9 | 9 | 9 | 8 |
| **Pc07** | 7 | 5 | 8 | 1 | 9 | 8 | 10 | 9 | 9 | 8 | 8 | 6 | 5 |
| **Pc08** | 3 | 5 | 7 | 1 | 8 | 5 | 10 | 10 | 10 | 8 | 8 | 7 | 7 |
| **Pc09** | 8 | 7 | 7 | 3 | 10 | 4 | 10 | 10 | 10 | 9 | 9 | 8 | 8 |
| **Pc10** | 9 | 2 | 7 | 1 | 10 | 9 | 10 | 9 | 9 | 7 | 7 | 5 | 2 |
| **Pc11** | 7 | 5 | 7 | 5 | 10 | 9 | 10 | 7 | 7 | 8 | 6 | 6 | 7 |
| **Pc12** | 7 | 8 | 8 | 1 | 8 | 7 | 10 | 8 | 8 | 6 | 8 | 5 | 8 |
| **Pc13** | 8 | 6 | 5 | 2 | 10 | 8 | 10 | 9 | 9 | 8 | 8 | 5 | 5 |
| **Pc14** | 8 | 7 | 7 | 5 | 8 | 8 | 10 | 10 | 10 | 9 | 8 | 7 | 8 |
| **Pc15** | 8 | 4 | 6 | 3 | 7 | 5 | 10 | 8 | 8 | 5 | 6 | 3 | 4 |
| **Pc16** | 7 | 6 | 6 | 3 | 8 | 8 | 10 | 9 | 9 | 8 | 5 | 4 | 3 |
| **Mean** | 7.7 | 5.7 | 6.6 | 3.0 | 8.9 | 7.4 | 9.9 | 8.9 | 8.9 | 7.7 | 7.3 | 5.9 | 5.8 |
| **SD** | 1.9 | 1.6 | 1.4 | 1.9 | 1.1 | 1.6 | 0.5 | 1.1 | 1.1 | 1.5 | 1.5 | 1.8 | 2.1 |

We asked participants: "for each type of terrain, please indicate how confident (or certain) you are of walking across it without losing balance, as though you had to step on it in real life outside. Please use a scale of 1 to 10, where 1 is not at all confident and 10 is extremely confident."
